# Neutrophil degranulation and extracellular ROS production are inactivated by *Yersinia pseudotuberculosis* YopE through a SKAP2 independent pathway

**DOI:** 10.64898/2026.06.02.729563

**Authors:** Pathricia A. Leus, Claudia Mañan Mejias, Alison Ren, Marjorie de la Rosa, Emma Lofgren, Steve C. Bunnell, David B. Sykes, Joan Mecsas

## Abstract

Upon sensing *Yersinia pseudotuberculosis (Yptb)*, receptor-mediated pathways are stimulated to trigger polymorphonuclear (PMN) antimicrobial responses. *Yptb* injects multiple Type 3 secreted effector proteins, Yops (*<u>Y</u>ersinia* <u>o</u>uter <u>p</u>roteins), that possess distinct biochemical functions, into PMNs to inhibit PMN responses. Here, we show that several Yops, YopE, YopH, and YopO, each partially interfered with CD63 mobilization to the plasma membrane, a marker for primary degranulation. The host pathways involved in CD63 mobilization are complex and it is not completely understood how Yops collaborate to inactivate this process. Here, CRISPR/Cas9 technology was used in an immortalized system of myeloid progenitor cells (Cas9-ER-HoxB8) to generate a panel of knockout PMN cell lines. To probe the impact of different Yops on the neutrophil pathways activated upon encountering *Yptb,* we interrogated the panel of genetically modified neutrophils with genetically modified bacteria. This approach of targeted gene deletion to inactivate specific pathways/proteins uncovered host pathways that synergize to induce CD63 mobilization that are distinctly targeted by YopE and YopH. YopE specifically inhibited CD63 mobilization in the absence of SKAP2, a YopH target, whereas YopH inhibited CD63 mobilization in the absence of RhoG, a YopE target, indicating that these Yops inactivate distinct signaling pathways contributing to CD63 mobilization. Furthermore, the SKAP2-independent pathway inactivated by YopE is involved in primary granule release and ROS production. Overall, this work highlights the diverse Yop-mediated mechanisms that WT-*Yptb* employs to effectively disarm PMN responses and provides an avenue to untangle neutrophil signaling pathways targeted by pathogens using Cas9-ER-HoxB8 cells.

**Author Summary:** When sensing invading bacteria, neutrophils become activated through multiple receptors that trigger signal-transduction cascades resulting in the generation antimicrobial responses. The enteric pathogen, *Yersinia pseudotuberculosis (Yptb),* is equipped to effectively inhibit these responses using its collection of effector proteins (Yops). Here, we developed a system to overcome the limitations of performing genetic manipulations in neutrophils by implementing CRISPR/Cas9 technology in an engineered system of myeloid progenitor cells (Cas9-ER-HoxB8) that can be induced to differentiate into neutrophils. By infecting genetically modified neutrophils with *Yptb* strains expressing individual Yops, we identified distinct host signaling pathways that synergize to induce neutrophil antimicrobial responses. Our findings provide insight into several signaling events triggered by *Yptb* infection and show how YopE and YopH target distinct pathways to block vesicle trafficking and extracellular ROS production. This powerful genetic system can be applied to other pathogens to dissect the intricacies of neutrophil-pathogen interactions.

## Introduction

Upon sensing the Gram-negative bacterium *Yersinia pseudotuberculosis (Yptb),* various neutrophil cell surface receptors are stimulated, including integrins, Toll Like Receptors (TLR), and G-protein-coupled-receptors (GPCRs) [1–4]. Activation of these receptors induces signaling cascades that trigger neutrophil effector responses including degranulation, extracellular vesicle (EV) release, phagocytosis, and reactive oxygen species (ROS) production. To counter these host defenses, the 3 human pathogenic *Yersinia* species: *Yptb, Y. enterocolitica,* and *Y. pestis,* inject 6-7 effector proteins, called Yops, into neutrophils via a Type 3 Secretion System (T3SS) [5, 6]. Neutrophils are the primary cell targets for Yop translocation during murine infection [7–9]. Collectively, Yops effectively inhibit many neutrophil effector responses, including degranulation [4, 7–12].

Neutrophils harbor different subsets of granules, primary, secondary, tertiary, and secretory, that are released in a sequential fashion upon sensing a pathogen in a process called degranulation. Primary granules are the last subset of granules released by neutrophils and contain the most potent antimicrobial molecules, including myeloperoxidase (MPO) and elastase. After *Y. pestis* infection, a combination of several Yops, YopE, YopH, or YopO (YpkA), coordinate to block CD63 plasma membrane localization, a marker for primary degranulation [11, 12]. In the context of *Yptb* infection, deletion of both YopE and YopH (Δ*yopEH)* abolishes *Yptb*’s ability to inhibit primary degranulation as measured by release of MPO, suggesting that one or both of these Yops are required to inhibit primary degranulation [10]. The requirement for multiple Yops to inhibit primary degranulation suggests that multiple pathways must be inactivated to completely block this process. This is supported by inhibitor studies with *Y. pestis,* which implicate both Ca^2+^ influx and Rac activity as the key processes targeted by YopE and YopH to inhibit primary granule mobilization [11]. However, the receptors and subsequent signaling events stimulated by the enteric *Yersinia* pathogens are distinct from those triggered by *Y. pestis*, with the enteric *Yptb* triggering integrin receptors, CCR5, and TLR-4, while *Y. pestis* triggers FPR1 [1–3, 13].

The biochemical activities of the Yops are conserved among the three *Yersinia* human pathogens. YopH is a tyrosine phosphatase and various proteins involved in integrin mediated neutrophil signaling, including Src Kinase Associated Phosphoprotein 2 (SKAP2), SH2 domain-containing leukocyte protein of 76 kDa (SLP-76), Vav guanine nucleotide exchange factor 1 (VAV1), and Phospholipase C Gamma 2 (PLCγ2) are dephosphorylated in the presence of YopH [4, 14–17]. Of note, the YopH target, SKAP2, is a key adaptor protein in neutrophil signaling and is required for maximal ROS production but not degranulation [4, 14]. Further, growth of a Δ*yopH* mutant is partially restored in a *Skap2^-/-^* mice, indicating that SKAP2 is one but not the only critical target YopH, and therefore other antimicrobial processes, such as degranulation, are involved in controlling the growth of Δ*yopH* [4]. YopE is a GTPase activating protein (GAP) and inactivates the small Rho GTPases, Rac1, Rac2, RhoA, and RhoG [18–22]. The YopE targets, Rac2 and RhoA, have both been implicated in neutrophil primary degranulation because of their involvement in actin cytoskeletal rearrangements. Specifically, activated Rac2 is essential for CD63 plasma membrane localization and actin polymerization [23–25], while inactivation of RhoA around the granules leads to blockage of actin polymerization, and thus allows for trafficking of the granule to its target membrane [26, 27]. To our knowledge before this work, RhoG had not yet been implicated in degranulation.

In this work, we developed a genetic system in neutrophils to identify specific proteins and signaling pathways targeted by different *Yptb* Yops to inhibit primary degranulation. Prior to this work, the ability to perform experiments with neutrophils containing targeted genetic deletions was limited. Previous approaches involve the use of bone marrow derived neutrophils from knockout (KO) or conditional KO mice and/or chemical inhibitors to inactivate the function of targeted proteins [4, 10, 11]. However, chemical inhibitors can have toxic, pleiotropic, and off-target effects, and PMNs isolated from KO mice can be limiting since targeting of some genes results in lethality, including *Rac1*, *RhoA*, and *Syk* [28–30]. While tissue conditional KOs have become more available, these are still limited and can be costly [31, 32].

To bypass these limitations and evaluate the roles of specific PMN genes, we generated a Cas9 expressing myeloid progenitor cell line using the conditional expression of HoxB8 under the control of estrogen (the ER-HoxB8 system) [33, 34]. This permitted targeted genetic deletions of many of the known and hypothesized Yop targets in immortalized ER-HoxB8 myeloid progenitor cells, which we then differentiated into PMNs (KO-PMNs). To examine the contribution of each Yop, we paired genetically modified host and genetically modified bacteria, where the newly generated KO-PMNs were infected with a panel of *Yersinia* strains expressing individual Yops (“Yop-only”). We hypothesized that inactivation of both a Yop-targeted pathway using a Yop-only strain and another pathway through deletion of the PMN gene would delineate distinct pathways targeted by individual Yops. Using this strategy, we uncovered that YopE acts independently of the YopH-targeted pathway, where YopE more effectively blocked degranulation and extracellular ROS production in the absence of the YopH target, SKAP2. Additionally, YopE’s impact on CD63 mobilization was enhanced in the absence of SLP-76 and PLCγ2, which are two other proteins dephosphorylated in the presence of YopH. Furthermore, our findings demonstrated that YopH acts on the RhoG-independent pathway to halt CD63 plasma membrane localization, overall supporting the idea that YopE and YopH target distinct signaling pathways.

## Results

### Creation of neutrophil knockout cell lines using a Cas9-GFP-ER-HoxB8 myeloid progenitor cell line

To uncover distinct neutrophil pathways impacted by different Yops, we performed a dual-genetics screen and infected a panel of neutrophil knockout cell lines (KO-PMNs) with various *Yptb* Yop mutants, each expressing an individual Yop. This first required the generation of neutrophil KO cell lines lacking genes that encode for known/hypothesized *yop* targets. Because neutrophils are difficult to genetically manipulate, we implemented CRISPR/Cas9 in ER-HoxB8-transduced myeloid progenitor cells [33, 34]. To make targeted deletions in myeloid progenitor cells, we harvested bone marrow cells from transgenic Cas9-GFP-expressing C57BL6/J mice and immortalized them through transduction with an estrogen-regulated (ER) HoxB8 construct to generate Cas9-GFP-ER-HoxB8 (Fig. 1A) [34].

**Figure 1:**
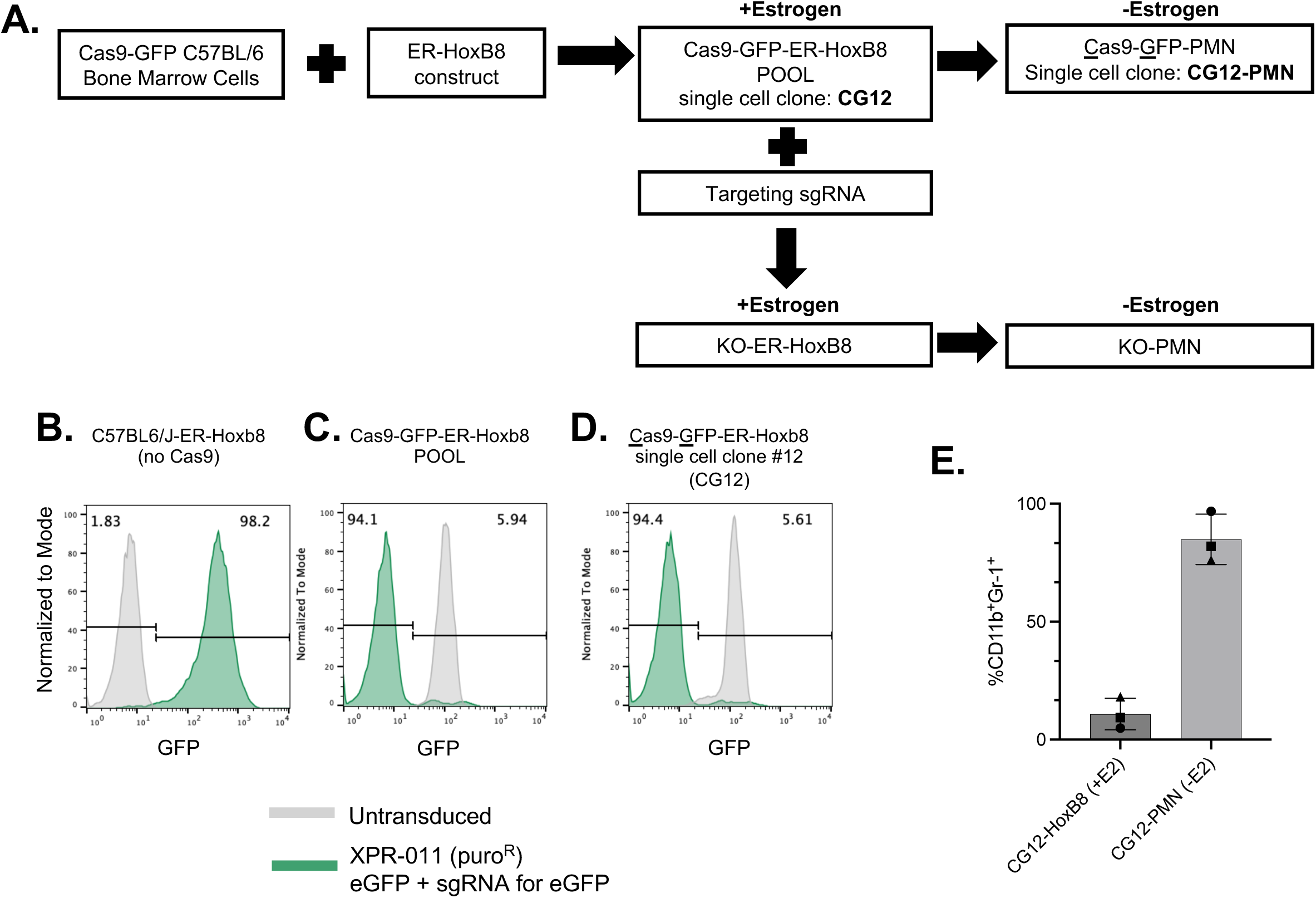
Cas9-ER-Hoxb8 cells have high Cas9 efficiency. (A) Schematic for generating knockout PMN (KO-PMN) cell lines. Myeloid progenitor cells isolated from bone marrow of a Cas9-GFP mouse was transduced with a construct encoding for an estrogen-regulated expression of HoxB8 to generate a pool of Cas9-GFP-expressing myeloid progenitor HoxB8 cells (Cas9-GFP-HoxB8s). <u>C</u>as9-<u>G</u>FP-ER-HoxB8 single cell clone, CG12, was transduced with sgRNAs to generate KO-ER-HoxB8 cells, which can be differentiated into KO-PMNs after removal of estrogen. (B-D) Histogram of (B) C57BL6/J-ER-HoxB8, (C) Cas9-GFP-ER-HoxB8 Pool, or (D) CG12, after transduction with pXPR-011 and 7 days of selection with 2 µg/mL puromycin. Cells were analyzed on BD LSRII for GFP expression (green peak) and compared to their respective untransduced cell line (gray peak). (E) CG12 was grown in the presence of estrogen (+E2) or was differentiated into CG12-PMN’s (-E2) (see Materials and Methods) for 3 days. CD11b and Gr-1 expression were assessed by Flow Cytometry. Each dot represents a biological replicate, which are matched by symbol. Bars are Mean ± SD of 3 biological replicates.

To assess the efficiency of Cas9 gene editing, we transduced the reporter construct, pXPR-011, into Cas9-GFP-ER-HoxB8s [35]. This construct contains (i) the gene encoding eGFP, (ii) a sgRNA for eGFP, and (iii) a puromycin-resistance cassette. In the presence of active Cas9, the sgRNA for eGFP will direct the Cas9 endonuclease to induce a double stranded break in both eGFP genes in the reporter construct and in the Cas9-GFP-ER-HoxB8 cells, resulting in loss of GFP expression. Thus, the absence of GFP expression is a proxy measure for Cas9 efficiency. ER-HoxB8 cells generated from WT C57BL6/J mice (WT-ER-HoxB8), which lack Cas9 and GFP, were used as a control and were successfully transduced, with 98.2% of the cells expressing GFP after puromycin selection (Fig. 1B). By contrast, 94.1% of the Cas9-GFP-ER-HoxB8 pool lacked GFP expression after transduction of pXPR-011, indicating that the Cas9 efficiently cleaved both the reporter GFP and the endogenous GFP in the cells (Fig. 1C). CG12, a single cell clone derived from the <u>C</u>as9-<u>G</u>FP-ER-HoxB8 pool, also exhibited robust Cas9 activity with 94.4% of cells losing GFP expression (Fig. 1D). This clone differentiated efficiently into CG12-PMN after removal of estrogen in the presence of IL-3 and GCSF (granulocyte colony stimulating factor), as demonstrated by increased expression of CD11b and Gr-1 (Fig. 1E) and was used as the parental strain for generating KO cell lines (see below).

### YopH, YopE, and YopO play roles in reducing CD63 plasma membrane localization after *Yptb* infection

To measure the impact of specific Yops in inactivating distinct neutrophil signaling pathways, we monitored CD63 movement to the plasma membrane after infection with *Yptb* mutants. CD63 is a transmembrane protein found on several vesicles, including primary granules and extracellular vesicles [36, 37]. Increased CD63 plasma membrane localization is widely used as a proxy to measure primary granule fusion with the plasma membrane [12, 36, 38]. To establish which Yops were involved in blocking CD63 plasma membrane localization in our neutrophil cell system, we infected CG12-PMNs with WT-*Yptb* and various *yop* mutants (Fig. 2). As expected, infection with a *Yptb-Δ5* mutant, which lacks the genes encoding for 5 Yop effectors (YopE, YopM, YopO, YopJ, and YopH) or a *Yptb* Δ6 mutant (a Δ5 mutant with subsequent deletion of the gene encoding for YopK) triggered high levels of CD63 plasma membrane localization in greater than 50% of the cell population (Fig. 2A, Fig. S1). Infection of CG12-PMN with WT-*Yptb,* however, resulted in comparable levels of CD63 plasma membrane localization to uninfected cells, indicating that WT-*Yptb* blocks CD63 plasma membrane localization effectively (Fig. 2A). Thus, ER-HoxB8-derived PMNs behaved like PMNs isolated from humans and mice, where one or more Yops block CD63 movement to the plasma membrane [10, 12, 38].

**Figure 2:**
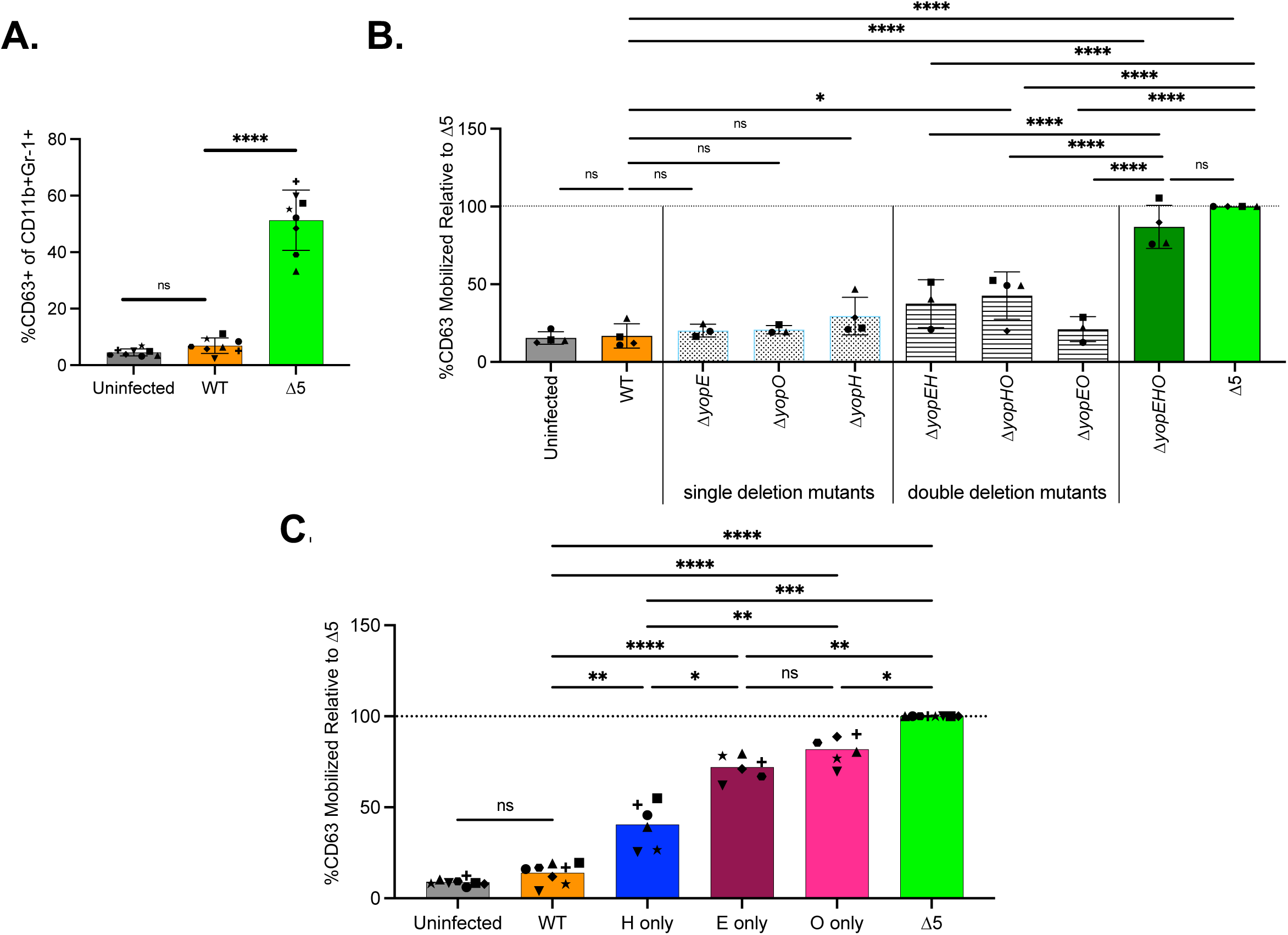
YopH, YopE, and YopO reduce CD63 plasma membrane localization after *Yptb* infection. (A-C) CG12-PMNs were infected with the indicated strains of *Yptb* at an MOI of 30 for 1 hour at 37°C. *Δ5* is a strain of *Yptb* lacking the 5 Yop effectors: YopE, YopH, YopO, YopJ, and YopM. After infection, cells were stained, fixed, and analyzed on Aurora Cytek Spectral Cytometer. Gating Strategy is shown in SF2. (A) Percentage of cells with CD63 localized to the plasma membrane are plotted. (B-C) Percentage of cells with CD63 localized to the plasma membrane plotted as a % of CD63 relative to Δ5, where Δ5 was set to 100%. Each dot is a biological replicate and each biological replicate represents an average of 2-3 technical replicates. Bars are the Mean ± SD of (A, C) 6-8 or (B) 3-5 biological replicates. Each dot represents a biological replicate, which are matched by symbol. Statistical significance was calculated using (A, C) One-Way ANOVA or (B) Mixed Effects Analysis followed by Tukey’s multiple comparison test. ns, not significant; P-value: *<0.05, **< 0.01, ***< 0.001, ****< 0.0001.

To further interrogate the roles of specific Yops, we infected CG12-PMNs with a series of single, double, and triple *yop* deletion mutant strains. Infection with a triple deletion mutant, Δ*yopEHO,* induced levels of CD63 plasma membrane localization comparable to *Yptb-Δ5* infected levels (Fig. 2B). This demonstrated that YopE, YopO, and/or YopH play roles in reducing mobilization of CD63-expressing vesicles to the surface and indicated that the combined activities of YopM and YopJ do not prevent CD63 movement (Fig. 2B). Infection of CG12-PMNs with Δ*yopE, ΔyopO,* or *ΔyopH* single deletion mutants resulted in blockage of CD63 plasma membrane localization comparable to infection with WT-*Yptb* (Fig. 2B). This result pointed to functional redundancies among YopH, YopE, and YopO. None of the double deletion mutants tested (Δ*yopEH, ΔyopHO,* or Δ*yopEO*) restored CD63 plasma membrane localization to *Yptb-Δ5* or Δ*yopEHO* infected levels (Fig. 2B). This data demonstrated that any of the 3 Yops are sufficient to block CD63 plasma membrane localization and that no single Yop is necessary for this blockage.

To assess if YopH, YopE, or YopO were sufficient to block or reduce CD63 movement to the plasma membrane after *Yptb* infection, we infected CG12-PMNs with strains of *Yersinia* expressing a single Yop from their native locus in a *Yptb-Δ5* background. These strains are referred to as “Yop-only” strains. Infection with “H-only” (Δ*yopEMOJ)* resulted in the most potent blockage of CD63 plasma membrane localization to approximately half of the CD63 mobilization observed in a *Yptb-*Δ5 infection (Fig. 2C). Infection with “E-only” (Δ*yopMOJH)* or “O-only” (Δ*yopEMJH)* resulted in modest but significant blockages of primary granule mobilization. Overall, no Yop was sufficient to block CD63 plasma membrane localization. Collectively, this data indicates that multiple overlapping pathways are involved in WT-*Yptb* blocking the trafficking of CD63-expressing vesicles to the plasma membrane and these pathways involve those targeted by YopH, YopE, and YopO.

### Generation of knockout cell lines in Cas9-GFP-ER-HoxB8 myeloid progenitor cells

The observation that YopH, YopE, and YopO, each partially reduced CD63 plasma membrane localization and that only a triple deletion strain of all three failed to suppress CD63 plasma membrane localization implies non-overlapping roles for each Yop. We hypothesized that inactivation of a Yop targeted pathway and inactivation of another pathway (targeted by a different Yop) through deletion of the targeted gene in PMNs could delineate distinct pathways targeted by the Yops. We used the CG12 clonal cell line to create an array of KO cell lines targeting genes that encoded for proteins that are known to be or potentially targeted by YopE, YopH, and YopO. These include small Rho-GTPases (YopE and YopO targets), tyrosine kinases and adaptor proteins (YopH and YopO targets), and several genes critical for antimicrobial functions of PMNs. CRISPR inactivation of the targeted genes appeared highly efficient, as observed with the CD63-KO cell lines, where transduction of two distinct sgRNAs targeting *Cd63* resulted in comparable levels of surface CD63 to unstained controls (Fig. S3A).

As a preliminary screen, the newly generated array of KO pools were infected with *Yptb-Δ5* and assessed for CD63 plasma membrane localization. Multiple KO cell lines showed reduced CD63 mobilization after *Yptb-Δ5* infection, the majority of which encode for proteins downstream of integrin receptor activation (Fig. S3B). Importantly, in most cases where genes were targeted with 2 or more sgRNAs, the generated KO pools phenocopied one another. To minimize the likelihood of any WT cells in the KO pools from outcompeting the KOs during passaging, cells from 13 of these pools were single cell cloned. The presence of a KO was confirmed using western blot, flow cytometry, and/or other functional assays (Fig. S4, S5) and the single cell closes were used for further analyses.

### Identifying key proteins involved in CD63 plasma membrane localization in response to *Yptb-Δ5*

To assess how the absence of the CRISPR-targeted genes impacted basal levels of CD63 plasma membrane localization, we measured the amount of surface localized CD63 in the uninfected panel of KO-PMN cell lines and compared it to the parental CG12-PMN cell line. An eGFP-KO single cell clone served as a control cell line that has undergone gene editing but should not impact any neutrophil function. In the absence of infection, there were lower levels of CD63 plasma membrane localization in many of the uninfected cells, including PLCγ2-KO, SYK-KO, SLP-76-KO, CYBB-KO, BTK-KO, and RhoG-KO (Fig. 3A). To evaluate whether the variable levels of CD63 on the surface of the cell were a result of changes in total amounts of CD63 protein in the KO cell lines, the levels of CD63 in permeabilized KO-PMNs were measured by flow cytometry. Total levels of CD63 were all comparable to or slightly higher than those in eGFP-KO or CG12-PMN; however, these levels were not statistically significant except for SLP-76-KO and SKAP2-KO, which were higher than the parental cell line (Fig. 3B). This result eliminated the possibility that lower basal levels of CD63 at the membrane resulted from lower overall levels of CD63 and suggests that these genes are all involved in normal membrane trafficking of CD63-containing vesicles to the plasma membrane.

**Figure 3:**
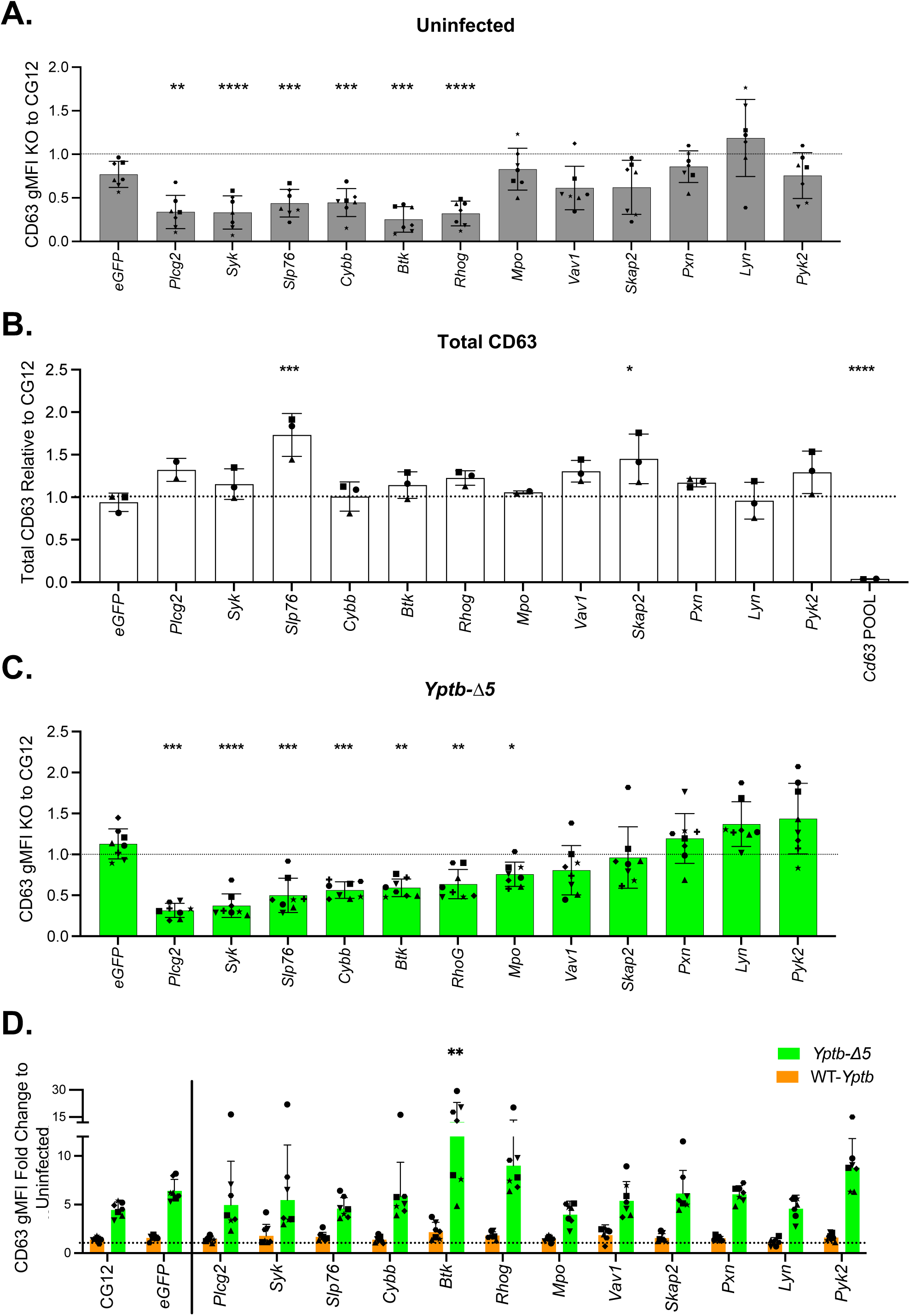
Identifying key proteins involved CD63 plasma membrane localization in response to *Yptb-Δ5*. (A-D) CG12-PMNs and single cell cloned KO-PMNs were (A, B, D) uninfected, (C, D) infected with *Yptb-Δ5 (*green) or (D) WT-*Yptb (*orange) with an MOI of 30 for 1 hour at 37°C. (A-D) Cells were stained, fixed, and analyzed on Aurora Cytek Spectral Cytometer. Gating Strategy is presented in SF2. CD63 geometric Mean Fluorescence Intensity (gMFI) of each knock out cell line was normalized as follows: (B) Cells were permeabilized prior to staining for total CD63 expression, (A, B) CD63 gMFI of uninfected KO cell line was divided by CD63 gMFI of uninfected CG12, (C) CD63 gMFI of the *Yptb-Δ5* infected KO cell line was divided by CD63 gMFI of the *Yptb-Δ5* infected CG12, (D) CD63 gMFI of infected conditions were normalized to uninfected CD63 gMFI of that same cell line to determine the fold change in CD63 surface expression. (A, C, D) n=7-8. (B) n=2-3. Statistical analysis was performed by comparing eGFP-KO with other KO cell lines using One-Way ANOVA followed by Dunnett’s test. Data are mean ± SD. Symbols are matched per biological replicate. P-values: *< 0.05; **< 0.01; *** < 0.001; **** <0.0001.

We next evaluated whether absence of the targeted gene in the single cell clones impacted CD63 movement to the plasma membrane after infection with *Yptb-*Δ5 (Fig. 3C). In this case, we compared the amount of CD63 on the cell surface of a knockout strain after infection with *Yptb-*Δ5 to the amount on CG12-PMN after infection with *Yptb-*Δ5. The absence of *Syk, Plcg2, Slp76,* and *Btk* demonstrated a significant defect in CD63 mobilization after infection with *Yptb-Δ5* compared to CG12-PMN and eGFP-KO (Fig. 3C), which is consistent with the fact that these proteins transmit signals downstream of β1 integrin activation. In addition, *Cybb* and *Rhog* also played important roles in regulating levels of CD63 on the membrane (Fig. 3C). These results were largely consistent with our observations of the pooled KO cell lines where CD63 plasma membrane levels were low after infection with *Yptb-*Δ5 (Fig. S3B). However, in the case of *Cybb,* the low levels observed in the single cell clone did not phenocopy the CYBB-KO pools generated from two different gRNAs or HoxB8-derived *Cybb^-/-^*PMNs generated from *Cybb*^-/-^ mice (Fig. 3A, 3C, Fig. S3B-C). Overall, this data identified key proteins involved in CD63 plasma membrane localization in response to infection with *Yptb-*Δ5, which includes RhoG, as well as key proteins that function downstream of integrins: PLCγ2, SYK, SLP76, and BTK (Fig. 3A, 3C, Fig. S3B-C).

Finally, we calculated the increase in CD63 at the plasma membrane after infection with *Yptb-*Δ5 compared to its basal level in each cell line (Fig. 3A) after infection with *Yptb-*Δ5 (Fig. 3D). Infection of CG12-PMN and eGFP-KO with *Yptb-Δ5* resulted in 4-6 fold increase in plasma membrane localized CD63 compared to their uninfected levels. With the exception of BTK-KO, infection of all the KO single cell clones with *Yptb-Δ5* resulted in a 4-10-fold increase of CD63 compared to their baseline uninfected levels (Fig. 3D). BTK-KO had a 12-fold increase which was significantly higher, although it is notable that the basal CD63 plasma membrane level was particularly low in BTK-KO (Fig. 3A). This data demonstrates that all KO cell lines were able to mobilize CD63 to comparable or greater levels as the control cell lines in response to infection with *Yptb-Δ5* (Fig. 3D). Moreover, the observation that knocking out these host proteins still resulted in mobilization of CD63+ vesicles after *Yptb-Δ5* infection points to the use of multiple host pathways involved in primary granule mobilization. Retention of some CD63 mobilization was critical for our next step in determining whether single Yops could better retard mobilization in specific KO cell lines. Importantly, WT-*Yptb* effectively blocked CD63 plasma membrane localization in all KO cell lines (Fig. 3D). This suggests that neutrophils lacking these specific proteins did not employ new or alternative pathways that could not be inhibited by one or more Yops.

### YopE and YopH target distinct signaling pathways to block CD63 plasma membrane localization

Based on the observation that individual Yop expression strains, “Yop-only” strains partially blocked CD63 mobilization, we reasoned that *Yptb-Δ*5 may induce CD63 mobilization through several distinct pathways upon stimulation of a variety of receptors (Fig. 4A). We next sought to identify PMN KOs that synergize with Yop-only strains in blocking CD63 plasma membrane localization relative to the Yop-only infected CG12-PMN. We hypothesized that we may observe enhanced Yop potency in blocking CD63 mobilization that could be due to inactivation of two distinct pathways: 1) a Yop-targeted pathway and 2) a second pathway inactivated by deletion of a PMN gene (Fig. 4A). Thus, we infected our KO cell lines with Yop-only strains. The impact of individual Yops on each KO cell line was quantified by calculating a ratio of CD63 plasma membrane localization in the Yop-only condition divided by the *Yptb-Δ5* infected condition. To compare the impact of the Yop in each KO cell line relative to the parental CG12-PMN, we divided the calculated ratio for each KO cell line by the ratio obtained from CG12-PMN (Fig. 4A). We referred to this normalized ratio as Relative Yop Potency, where a value less than 1 indicates enhanced potency of the Yop in the KO cell line compared to the parental CG12-PMN cell line. This would signify that the Yop-targeted pathway and the protein encoded by the knocked-out gene belong to distinct pathways. On the other hand, if the Yop has the same impact in the absence of the targeted gene compared to the parental CG12-PMN line, then the Relative Yop Potency should be around 1, indicating that the deletion of the PMN gene did not disrupt additional pathways to the ones targeted by the Yop. Such a result could arise because either the KO has no impact on CD63 mobilization or the KO is in the Yop targeted signal-transduction pathway.

**Figure 4:**
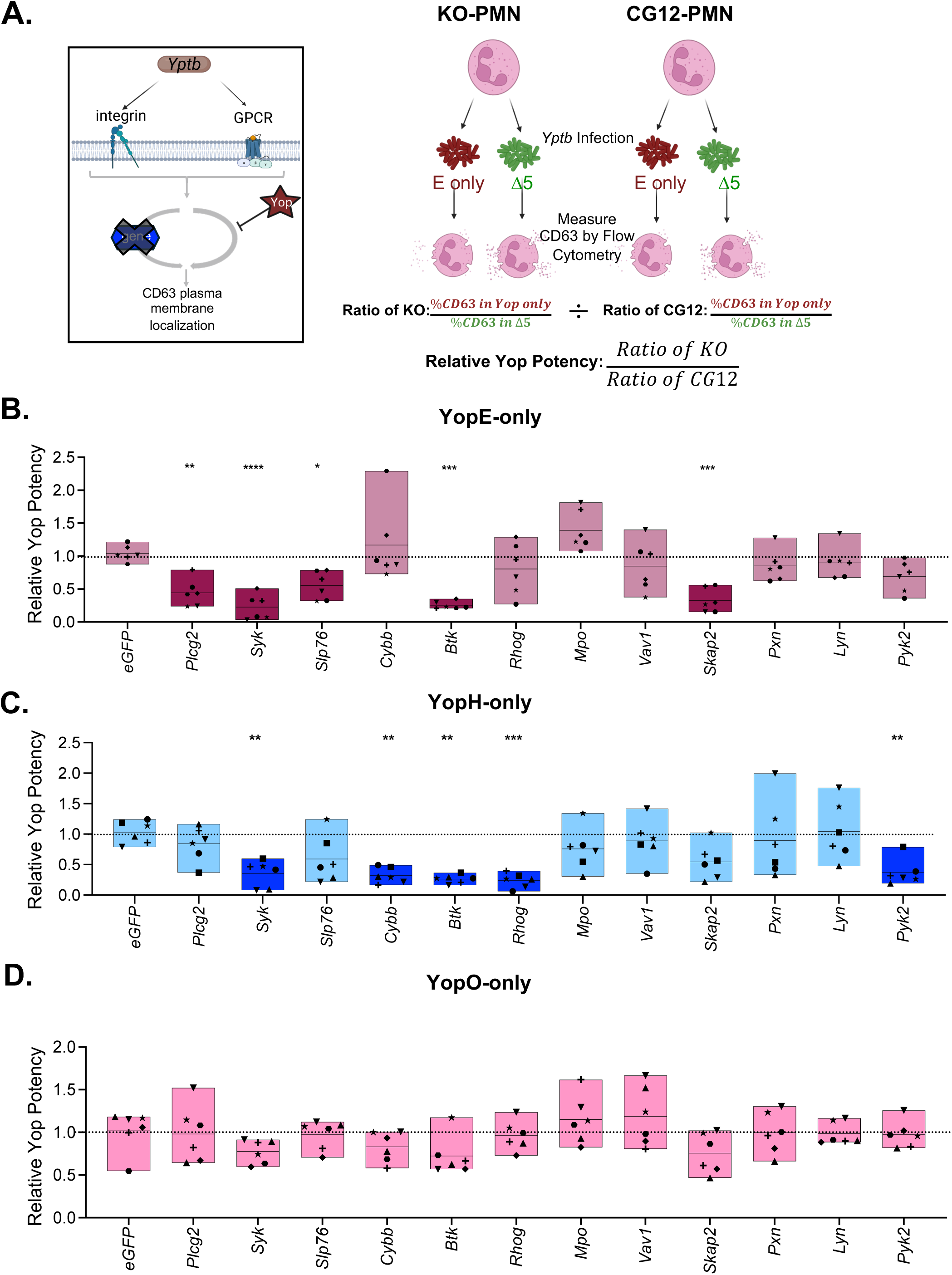
YopE and YopH target distinct signaling pathways to block CD63 plasma membrane localization. (A) Graphical representation of model (Left) and experimental design and set up for analysis (Right). For each KO cell line: Relative Yop Potency was calculated by dividing %CD63+ of Yop-only infected cells by %CD63+ of the Δ5-infected cell line. This ratio was divided by the same ratio obtained from the parental CG12-PMN. The normalized CD63 of KO to CG12 is plotted as Relative Yop Potency in (B-D). CG12, and KO cell lines were differentiated into KO-PMNs and infected with (B) YopE-only, (C) YopH-only, and (D) YopO-only with an MOI of 30 for 1 hour at 37°C. After infection, cells were stained, fixed, and analyzed on Aurora Cytek Spectral Cytometer. Cells were gated as shown in Fig. S1 and percentage of CD63+ cells of each knock out cell line was normalized. N=6 biological replicates are indicated by a symbol, each biological replicate had 2-6 technical replicates. Statistical analysis was performed by comparing eGFP-KO with KO cell lines using RM One-Way ANOVA followed by Dunnett’s test. Symbols are matched per biological replicate. Bars represent minimum to maximum and line indicates mean. P-values: *< 0.05; **< 0.01; *** < 0.001; **** <0.0001.

Infection of the eGFP-KO control with the Yop-only strains resulted in comparable levels of blockage to the CG12-PMN parental cell line (Fig. 4B-4D). Notably, infection of our panel of KOs with YopE-only revealed that YopE-only was significantly better at uniquely dampening CD63 plasma membrane localization in the absence of SKAP2, SLP-76, and PLCγ2 (Fig. 4B). This indicates that when these pathways are absent or disrupted, YopE’s ability to inactivate the signaling for CD63 mobilization becomes more apparent. We and others have previously shown that SKAP2, SLP-76 and PLCγ2 are dephosphorylated in the presence of YopH [4, 7, 17, 39]. This result suggests that YopE targets and inactivates a pathway that is independent of this YopH-targeted pathway, supporting the notion that YopH and YopE function on separate pathways to reduce CD63 plasma membrane localization. YopH-only was significantly better at uniquely dampening CD63 plasma membrane localization in the absence of RhoG, CYBB, and PYK2 (Fig. 4C). The small Rho GTPase, RhoG, is a known target of YopE [21, 22]. Thus, the observation that the impact of YopH-only was enhanced in the absence of RhoG indicates that YopH inactivates a signaling pathway that is independent to the YopE-mediated pathway [21, 22]. In contrast with YopE-only and YopH-only strains, infection with YopO-only did not further reduce CD63 plasma membrane localization in any of the KO-PMN cell lines. Additionally, both YopE-only and YopH-only were more effective at blocking the residual signaling for CD63 mobilization in the SYK-KO and BTK-KO, indicating that the impact of YopE and YopH is enhanced when the pathway propagated by these proteins is impaired.

### YopE mitigates primary granule release and extracellular ROS production in a SKAP2-independent manner

Since CD63 mobilization was relatively unimpaired after infection with *Yptb*-Δ5 in SKAP2-KO PMNs (Fig. 3C), we sought to further understand the impact of YopE on antimicrobial responses in this host background. To further validate that YopE targets a SKAP2-independent pathway to inhibit CD63 plasma membrane localization, we assessed the impact of YopE on CD63 mobilization in an independent SKAP2-KO clone generated with a different sgRNA, F8-4. YopE-only had similar impacts on both SKAP2-KO single cell clones (Fig. 5A). Since CD63 plasma membrane localization serves as a proxy for primary degranulation, we measured release of the primary granule component myeloperoxidase (MPO) in the supernatant of infected cells to directly measure primary granule release. WT-*Yptb* blocked MPO release to levels seen in uninfected CG12-PMN and SKAP2-KO, while *Yptb-Δ5* infection resulted in a 2-fold increase compared to uninfected (Fig. 5B). In CG12-PMN, infection with YopE-only reduced MPO release but was not sufficient to completely block MPO release to uninfected levels. However, in the absence of SKAP2, infection with YopE-only blocked MPO release to the same degree as the WT-*Yptb* condition (Fig. 5B). These results phenocopy the CD63 results and indicate that YopE-only inhibits primary granule mobilization and exocytosis in the absence of SKAP2. By contrast, infection with YopH-only effectively blocked MPO release, but only reduced CD63 plasma membrane localization to about half of what was observed in the *Yptb-Δ5* infected CG12-PMN (Fig. 2C, Fig. 5B). This indicates that while YopH is sufficient to block release of primary granules completely, some CD63+ vesicles can still mobilize to the plasma membrane.

**Figure 5:**
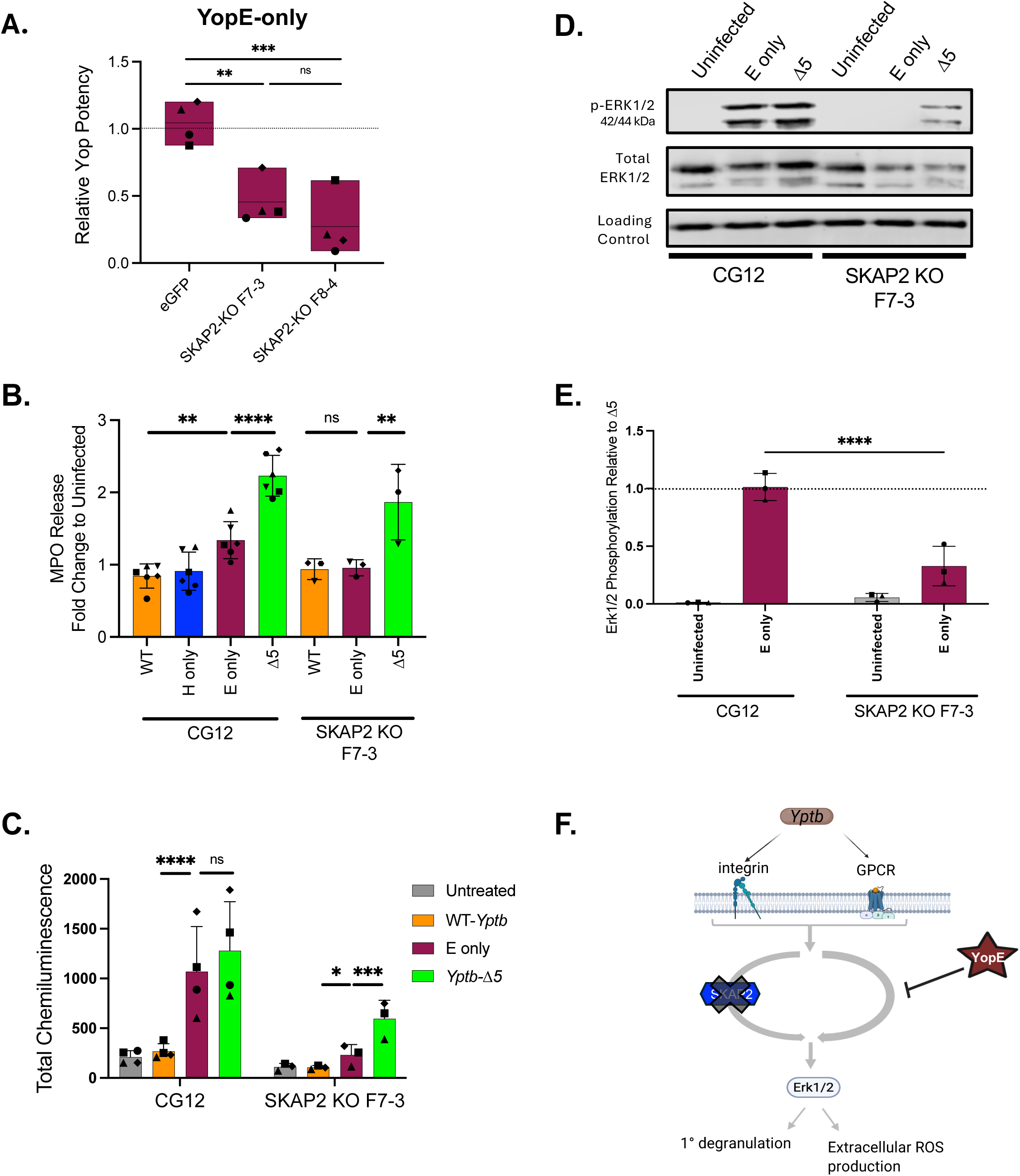
YopE mitigates primary granule release and extracellular ROS production in a SKAP2-independent manner. (A-E) CG12, eGFP-KO, SKAP2-KO F7-3, and (A) SKAP2-KO F8-4 were differentiated into PMNs. (A) Hoxb8-PMNs were infected with *Yptb* YopE-only and *Yptb-*Δ5 at an MOI of 30 for 1 hour. Cells were stained, fixed, and analyzed on Aurora Cytek Spectral Cytometer. Cells were gated as shown in Supp F1. CD63 gMFI of each knockout cell line was normalized as detailed in Fig. 4A. Statistical analysis was performed against eGFP using One-Way ANOVA followed by Dunnett’s multiple comparisons test. (B-E) Cells were infected with the indicated *Yptb* strains at an MOI of 20 for (B) 1 hour, (C) 30 min, and (D-E) 5 min. (B) After 1 hour of infection, supernatant was collected to quantify MPO released via ELISA. The concentration of MPO in each sample was divided by the concentration detected in the uninfected condition of each respective cell line and reported as fold-change. The average of 3-6 biological replicates, with 2 technical replicates is plotted. Statistical significance was determined using Mixed-Effects Analysis followed by Dunnett’s test against E-only condition. (C) Extracellular ROS production was measured via an isoluminol chemiluminescence assay. Chemiluminescence was recorded every 2 min for 30 min of infection and area under the curve (AUC) was calculated and normalized according to cell number using Cell Titer Glo. AUC values were log transformed and statistical significance was calculated using Two-Way ANOVA followed by Tukey’s multiple comparison test. (B-C; D) each symbol represents one biological replicate with same symbol indicating samples were collected on the same day. (D-E) Lysates were immunoblotted with anti-phospho-Erk1/2; anti-RhoGDI was used as a loading control for all blots. Phospho-blots were stripped with 7M Guanidine HCl and probed with anti-Erk1/2 as a control. (D) Representative blot of quantified biological replicates in (E). (E) p-ERK1/2 signal for each biological replicate of was normalized to RhoGDI for each experiment. The values from uninfected and E only were then divided by the value from *Yptb-Δ5* for each respective cell line. Normalized Erk1/2 phosphorylation relative to Δ5 are plotted. n=3. Statistical significance was determined using Two-Way ANOVA followed by Sidak’s multiple comparison test. (F) Summary model. P-values: *< 0.05; **< 0.01; *** < 0.001; **** <0.0001.

Data from our lab have previously demonstrated that SKAP2 is required for maximal extracellular ROS production upon stimulation of integrin and GPCRs [4, 40]. Here, we assessed whether YopE blocks extracellular ROS in the absence of SKAP2 after infection with WT-*Yptb*, YopE-only or *Yptb-*Δ5. SKAP2 was required for maximal extracellular ROS production after *Yptb-Δ5* infection, but modest amounts of extracellular ROS were still detected in the absence of SKAP2 (Fig. 5C). YopE-only did not block ROS production in CG12-PMN as ROS levels were comparable to *Yptb*-Δ5 (Fig. 5C). However, in the absence of SKAP2, infection with YopE-only significantly reduced the remaining extracellular ROS produced (Fig. 5C). Collectively, these data suggest that YopE mitigates primary granule release and extracellular ROS in a SKAP2-independent manner.

To evaluate the impact of YopE in the SKAP2-independent pathway relative to distal signaling events, we assessed activation of Extracellular Regulated Kinase 1/2 (ERK1/2). In WT-PMNs, SKAP2 is important for ERK1/2 activation, which leads to p47^phox^ translocation, activation of the NADPH oxidase complex, and ROS production [4, 41, 42]. In the absence of SKAP2, modest amounts of ERK1/2 activation were observed after infection with *Yptb-*Δ5 (Fig. 5D-E) [4]. In a SKAP2-KO, infection with a YopE-only strain blocked ERK activation, demonstrating that the inhibitory activity of YopE occurs upstream of ERK activation (Fig. 5D-E). These results are consistent with the ROS data, where the modest amounts of extracellular ROS produced in a *Yptb-Δ*5 infected SKAP2-KO was reduced by YopE. Overall, this indicates that YopE reduces ERK activation via a SKAP2-independent signal-transduction pathway that occurs upstream of ERK1/2 (Fig. 5F).

### Δ*yopE* is not restored for growth in *Skap2-/-* mice

Since our findings demonstrate that in the absence of SKAP2, YopE inhibits PMN primary granule release, CD63 plasma membrane mobilization, and extracellular ROS production, we predicted that a Δ*yopE* mutant strain should not be restored for growth in a *Skap2*^-/-^ mouse. On the other hand, if YopE inactivates a pathway that requires SKAP2, we would expect that a Δ*yopE* mutant would be more competitive in a *Skap2*^-/-^ mouse and thus have a higher competitive index than it does in a BALB/c WT mouse. This result can be compared to previously published competition experiments with Δ*yopH,* which demonstrated that Δ*yopH* colonization is partially restored in the absence of *Skap2,* indicating that YopH targets a SKAP2-dependent process [4]. To test this, BALB/c WT and *Skap2^-/-^*mice were orally (Fig. 6A) or intravenously (Fig. 6B) infected with equal doses of WT-*Yptb* (kan^R^) and *ΔyopE.* As expected, Δ*yopE* had a defect in colonizing BALB/c WT small intestinal contents upon oral infection (Fig. 6A) [20]. While there was a slight trend towards *ΔyopE* being more competitive in the *Skap2*^-/-^mouse, it was not significant and the same trend held when the *Skap2*^-/-^ mouse was challenged with the isogenic WT-*Yptb* (kan^S^) versus the WT-*Yptb* (kan^R^). Intravenous administration of *Yptb* similarly demonstrated that Δ*yopE* was defective in colonizing the spleen in the systemic infection model in both the BALB/c WT and *Skap2*^-/-^ mice (Fig. 6B). Combined, these data support our observation in the ER-HoxB8 system in which YopE targets a SKAP2-independent pathway.

**Figure 6:**
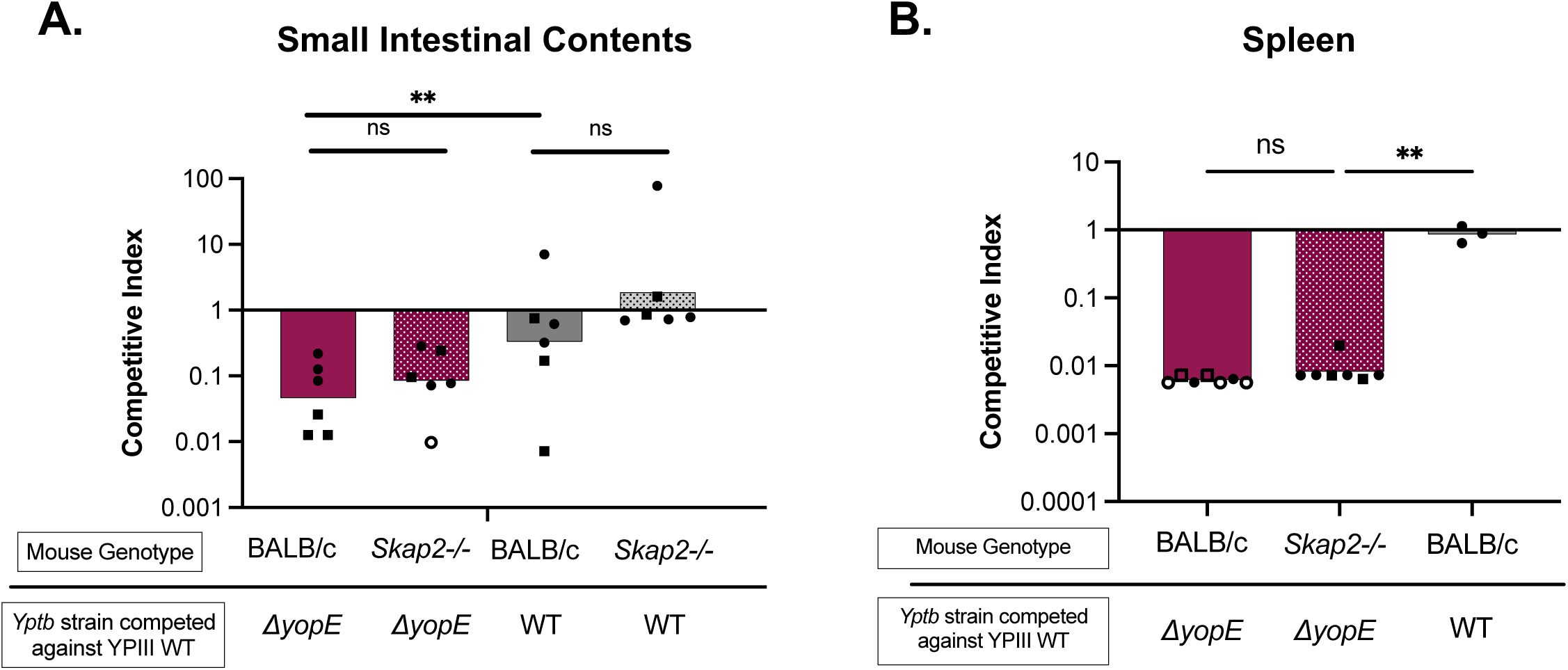
Δ*yopE* is not restored for growth in *Skap2^-/-^* mice. (A-B) BALB/c WT and BALB/c *Skap2^-/-^* mice were (A) orally infected with equal doses of *Yptb* YPIII (kan^R^) and Δ*yopE* for a total CFU of 2 x 10^9^ or (B) intravenously infected with equal doses of *Yptb* YPIII (kan^R^) and Δ*yopE* for a total CFU of 1 x 10^3.^ After 3 days of infection, (A) small intestinal contents or (B) spleen were harvested and homogenized and outputs were plated on L plates containing irgasan. Outputs were then patched on L plates containing kanamycin to determine the competitive index (CI). Each dot represents a mouse. Open symbols indicate only WT^kan^ was detected in 100 colonies patched. CI values were log transformed and statistical significance was calculated using One-Way ANOVA followed by Dunnett’s comparison test. P-value **< 0.01.

## Discussion

In this work, we developed and utilized a powerful system to generate genetic deletions in neutrophils and deconvolute the contributions of YopE and YopH in host neutrophil signaling pathways that converge on CD63 mobilization, a key marker of primary degranulation. This robust Cas9-ER-HoxB8 system can be used in screens to study neutrophil biology and applied to probe interactions of pathogens with neutrophils. Overall, our findings indicate that multiple host pathways are triggered by *Yptb-*Δ5 to induce CD63 mobilization and primary granule mobilization. Despite this multifaceted response to infection by *Yptb-*Δ5, WT-*Yptb* effectively blocked CD63 plasma membrane localization, demonstrating that *Yptb* is adeptly armed to halt CD63 mobilization. In HoxB8-PMNs, the absence of one or two of YopH, YopE, and YopO still resulted in significant blockage of CD63 mobilization and injection of YopH-only, YopE-only or YopO-only partially blocked CD63 mobilization, indicating each Yop plays a role in preventing CD63 mobilization. On the host end, we identified RhoG and confirmed that PLCγ2, SYK, SLP76, and BTK all play central roles in the process of CD63 mobilization in response to *Yptb- Δ5* infection (Fig. 7). Our findings also demonstrated that YopE’s contributions to blocking CD63 mobilization, primary granule release, and extracellular ROS production occur in a SKAP2-independent manner. Furthermore, YopE effectively prevented the residual signaling that remained in the absence of key proteins involved in integrin-mediated signaling (PLCγ2, SYK, SLP76, BTK) suggesting that YopE can block CD63 mobilization that is triggered by a non-integrin receptor.

**Figure 7:**
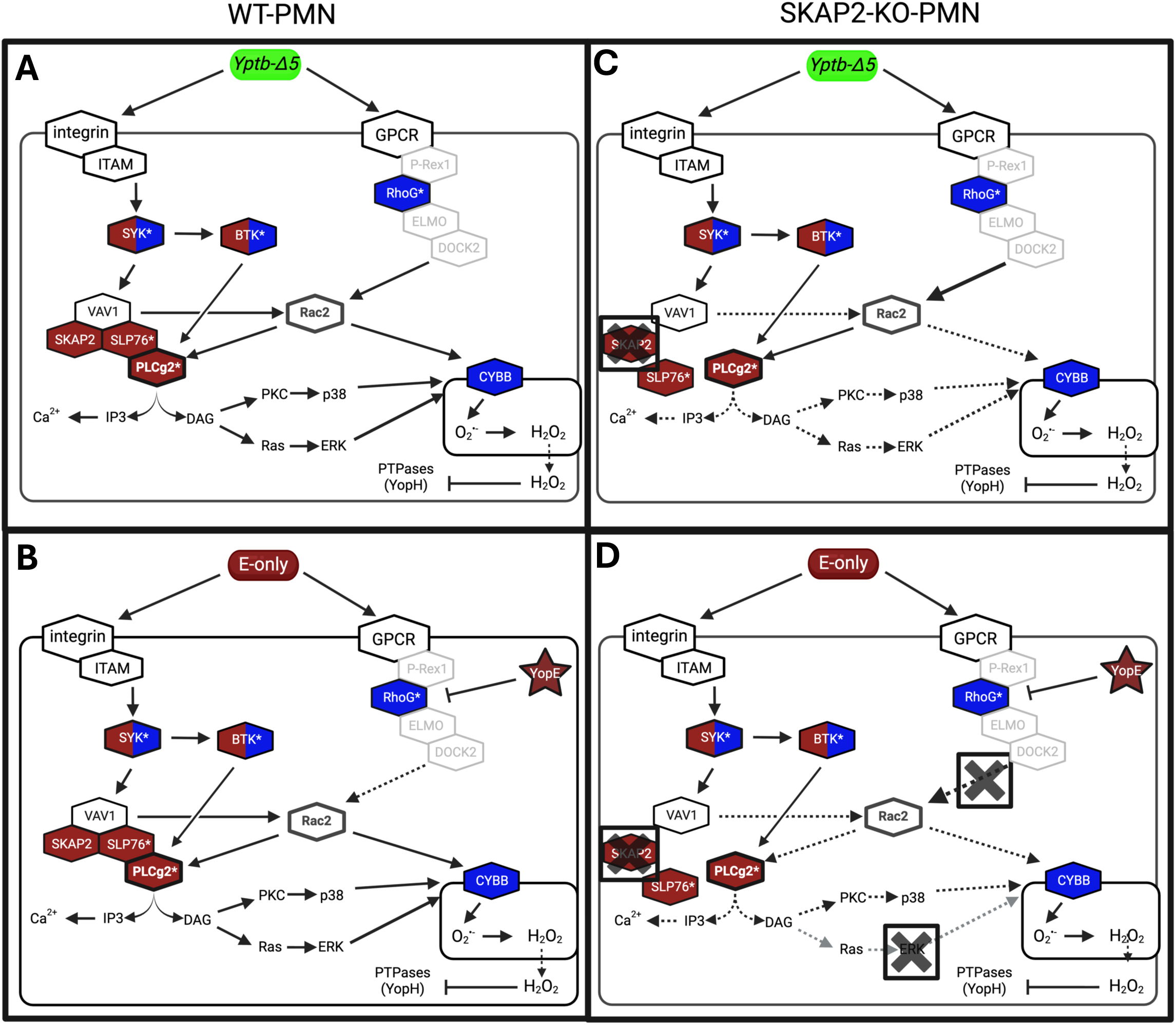
Proposed Model. Our work identified proteins that play a critical role in CD63 plasma membrane localization upon *Yptb* infection (denoted with *). Two proteins in particular, Rac2 and PLCγ2 (thicker borders), play an important role in actin cytoskeletal rearrangements and Ca^2+^ flux, respectively, and are required for primary degranulation. During integrin outside-in signaling in WT-PMNs infected with *Yptb Δ5* (A), ITAM bearing adaptors initiate downstream pathways involving recruitment and activation of SYK, leading to phosphorylation of BTK, as well as formation of a signaling complex involving VAV1, SKAP2, SLP-76, and PLCγ2, which YopH can inactivate. PLCγ2 cleaves PIP2 into IP3 and DAG which can each induce Ca^2+^ flux and downstream activation of p47phox and subsequent activation of the NADPH oxidase complex via PKC-p38 and Ras-ERK pathway. The signaling complex containing VAV1 is involved in activating Rac2, which is critical for primary degranulation and ROS production. Another mechanism for Rac2 activation is likely through a GPCR-mediated pathway driven by RhoG. RhoG is activated by the GEF, P-Rex1, and upon activation can recruit ELMO/DOCK2, which functions as a GEF to activate Rac2. YopE (star symbol) can directly inactivate RhoG but only has a modest impact on blocking both primary degranulation and ROS in WT-PMNs (B). In the absence of the scaffolding protein SKAP2 (C-D), the protein complex with VAV1, SLP76, and PLCγ2 is impaired, and VAV1 can no longer activate Rac2. Thus, the activation of Rac2 becomes completely reliant on GPCR-signaling via RhoG. In a SKAP2-KO, YopE can directly inactivate RhoG (D), subsequently impeding activation of Rac2 and effectively inhibiting both primary degranulation and extracellular ROS production. In the absence of proteins colored in maroon, YopE-only was more potent in blocking CD63 plasma membrane localization. In the absence of proteins colored in blue, YopH-only was more potent in blocking CD63 plasma membrane localization. YopE and YopH were both more potent in the absence of proteins colored in both blue and maroon. Proteins outlined in grey were not tested in this work.

YopE functions as a GAP for the small Rho GTPases RhoA, Rac1, Rac2, and RhoG [18–21, 43]. In neutrophils, several small Rho GTPases tune actin cytoskeletal dynamics to permit primary granule mobilization [11, 23–26]. Activation of Rac2 is required for CD63 plasma membrane localization [24]. By contrast, RhoA inactivation is critical for granule translocation, which occurs by the host encoded GAP, GMIP, and results in inhibition of F-actin assembly around the granules, thus facilitating granule translocation to the plasma membrane [26]. Here, infection with YopE-only had a modest impact in blocking CD63 plasma membrane localization in the control cell lines CG12-PMN or eGFP-KO. Since Rac2 activation is essential for primary granule release [23, 25], YopE’s modest blockage of CD63 in CG12-PMN suggests that Rac2 is not a direct or the primary target in these parental cells. Notably, the inhibitory effect of YopE on CD63 mobilization and primary granule release was more pronounced in the absence of SKAP2. This indicates that in the absence of SKAP2, YopE either more efficiently targets Rac2, or more likely, that YopE is effectively targeting an upstream small Rho GTPase that leads to Rac2 activation.

We postulate that there are at least two distinct pathways driving the activation of Rac2 (Fig. 7). In WT-PMNs infected with *Yptb-*Δ5, Rac2 is activated by VAV1/3 through a SKAP2-mediated mechanism, likely through its scaffolding function (Fig. 7). The absence of SKAP2 reveals the importance of a second pathway to activate Rac2, which we suspect is driven by RhoG via a GPCR-mediated pathway (Fig. 7). A role for RhoG is supported by our findings that the absence of RhoG resulted in a defect in steady state levels of surface-localized CD63 as well as on CD63 mobilization upon infection with *Yptb-Δ5* (Fig. 3). Damoulakis et al. have shown that there are RhoG-dependent and RhoG-independent mechanisms to activate Rac2 [44].

Additionally, a previous study showed that Rac activation in neutrophils is severely impaired when the GEFs, VAV1/3 and P-Rex1, are all deleted compared to individual deletions of all members of the VAV family or P-Rex1, highlighting the two distinct pathways to activate Rac1/2 [45]. P-Rex1 is a known GEF for RhoG and can activate Rac2 via ELMO/DOCK2 [44, 46]. The fact that we observed no enhanced impact of YopE in the absence of VAV1 is likely because there is redundancy with VAV3, another member of the VAV family [47]. In addition, while there were reduced levels of CD63 mobilization in the absence of BTK, PLCγ2, SYK, or SLP76, YopE was effective at blocking the residual signaling in these KO cell lines. Thus, we propose that when the signaling propagated by the protein microcluster formed by SKAP2-SLP-76-PLCγ2 is damaged genetically, the signaling for Rac2 activation becomes reliant on the RhoG-mediated pathway that may be triggered through activation of one or more GPCRs that sense bacteria, and YopE impedes CD63 mobilization, primary degranulation, and ROS production by inactivation of RhoG (Fig. 7).

YopH was the most potent single Yop at inhibiting CD63 mobilization and was completely effective at blocking MPO release. YopH inactivates host pathways governing Ca^2+^ influx and actin cytoskeletal rearrangements, both of which are important processes mediating primary degranulation [16]. By inactivating PLCγ2, YopH inhibits cleavage of PIP_2_ into IP_3_ and DAG, effectively inhibiting Ca^2+^ influx and activation of extracellular ROS production, respectively [4, 14]. YopH also targets the signaling complex involving VAV1 [14] for inactivation, thereby preventing activation of Rac1/2 and inhibiting NADPH oxidase activation and actin cytoskeletal rearrangements required to traffic the granules to the plasma membrane. The potency of YopH is enhanced in the absence of RhoG, CYBB, and PYK2. One intriguing common phenotype of CYBB, RhoG, and PYK2 is that they all play roles in ROS generation [42, 48, 49]. The possible link between generation of ROS and CD63 mobilization after *Yptb* infection remains to be explored experimentally. Studies in T cells have demonstrated that ROS can function as a signaling molecule to regulate cell signaling through transient inactivation of PTPases [50, 51]. Thus, it is tantalizing to speculate that in the absence of host generated ROS through genetic deletion of CYBB, RhoG, or PYK2, inactivation of host PTPases and YopH is reduced and thus more potent inhibition of tyrosine kinases signal transduction relays occurs by PTPases.

While YopH-only was sufficient to block MPO release, it did not block CD63 plasma membrane localization to the same levels as WT-*Yptb.* YopH’s enhanced potency in blocking CD63 mobilization in the absence of SYK, CYBB, BTK, RhoG, or PYK2 indicates that YopH targets pathways independent of these proteins and these proteins are part of signal-transduction pathways that contribute to CD63 mobilization after *Yptb* infection. *Y. pestis* was recently shown to block release of neutrophil-derived EVs, which are enriched with CD63 [52, 53]. In particular, YopH, along with YopE and YopK, played a role in altering proteins packaged in EVs [52]. It is possible that YopH’s enhanced potency in reducing CD63 plasma membrane localization points to the involvement of these host proteins in regulating processes involved in CD63+ vesicle trafficking or uptake of EVs. EV biogenesis, uptake, and release of primary granules all involve vesicle trafficking and are at some level interconnected but the mechanistic details and roles of these host proteins in these processes require further investigation.

Our work and others have demonstrated that SYK and BTK are critical for CD63 mobilization, yet residual CD63 mobilization occurs after infection with *Yptb-Δ5* [54–56]. SYK is a proximal protein downstream of integrin [56] and can recruit and phosphorylate BTK. Studies have shown that BTK activates the YopH target PLCγ2 [57, 58], and that BTK functions upstream of Rac2 [54]. In this work, we found that both YopH-only and YopE-only infection led to enhanced blockage of CD63 mobilization the absence of BTK and SYK. In the case of YopE, we suspect that SYK and BTK are not required to propagate the RhoG-mediated signaling, therefore allowing for enhanced potency of YopE in this parallel or independent pathway (Fig 7). On the other hand, in the SYK-KO and BTK-KO, the residual and impaired signaling mediated by the SKAP2-SLP-76-VAV1-PLCγ2 axis is effectively blocked by YopH, making YopH’s inhibitory effect more apparent.

Our observation that YopH, YopE, and YopO all contribute to blocking CD63 mobilization in ER-HoxB8 PMNs is largely consistent with previously published results in human and murine neutrophils, which showed that YopE and YopH were involved in blocking degranulation [4, 10–12]. In addition to YopE and YopH, Pulsifer et al. also reported a role for YopJ and YopK, which we did not observe in our system [12]. Some possible explanations for the discrepancy could be a result of differences in the relative abundance of signal transduction proteins between human and murine derived PMNs or differences in receptors triggered between *Yptb* and *Y. pestis*.

In conclusion, the use of our dual-genetics approach demonstrates that there are at least two distinct host signaling responses which synergize to mobilize CD63 to the plasma membrane, one targeted by YopH and one by YopE. However, our panel of targeted genes did not provide additional insight into YopO’s targeted pathways, and a focus of our future studies is to uncover how YopO is contributing to this process. This work provides significant mechanistic insight into and untangles some of the complex receptor-mediated neutrophil signaling triggered upon infection with *Yptb.* Ultimately, we demonstrated how YopE and YopH individually contribute to inactivate host signaling to block CD63 mobilization and ROS production. Finally, we developed a powerful system to generate genetic deletions in neutrophils, a key innate immune cell-type that is notoriously difficult to manipulate genetically.

## Materials and Methods

### Generation of Cas9-GFP-ER-HoxB8’s

ER-HoxB8 cells were generated according to previously described protocols with the following modifications [34, 40]. To generate Cas9-GFP expressing ER-HoxB8 cells, the Spinflush method was used to harvest bone marrow cells from the femurs and tibias of 10-week-old C57BL/6J female mice (Rosa26-Cas9 knockin on B6J) from The Jackson Laboratory (Strain #26179). Briefly, one end of the bone was cut to expose the marrow and placed faced down into a microcentrifuge tube. Bones in the microcentrifuge tube were spun at 10,000 x g for 30 seconds at room temperature to flush the marrow. Bone marrow cells were resuspended in flow cytometry buffer and passed through a 40 µm filter. Cells were then carefully layered over Lympholyte M (Cat# CL5031) and centrifuged at 1000 x g for 20 minutes at room temperature. The cell suspension was then isolated, spun, and resuspended in RPMI + 10% v/v murine stem cell factor (mSCF) secreted from Chinese hamster ovary (CHO) cells + 10 ng/mL IL-3 + 10 ng/mL IL-6 and cultured for 24-48 hours prior to transduction.

The ER-HoxB8 construct, MSCV-neo-HA-ER-Hoxb8, was packaged in a lentivirus [34]. For transduction, bone marrow cells were seeded on a fibronectin-coated 12 well plate at 5 x 10^5^ per well and transduced via Spinoculation in the presence of 8 µg/mL polybrene at 1000 x g for 90 minutes at room temperature. ER-HoxB8 cells were grown in GMP Media: complete RPMI (RPMI-1640, 10% FBS, 100 U/mL Penicillin and 0.1 mg/mL streptomycin) supplemented with 0.5 µM β-estradiol and 1.25% v/v mSCF. Successfully transduced cells with ER-HoxB8 were selected using 1mg/mL G418. Cell lines derived from single cell clones were generated by limiting dilution and individual clones were screened for low CD11b baseline expression in undifferentiated cells. To ensure that Cas9 was functional in Cas9-GFP-ER-HoxB8 clones, cell lines were screened for Cas9 activity with a reporter plasmid (pXPR-011; [35]). Cas9-GFP single cell lines were also assessed by flow cytometry for appropriate differentiation through surface expression of Gr-1 and CD11b or through cell cycle analysis with DyeCycle (Invitrogen Cat# V35003) after removal of estrogen in the media. Single cell line, CG12, was selected for the parental cell line for the generation of KO cells.

### Differentiation of ER-HoxB8 cell lines into neutrophils

ER-HoxB8 cells were washed 3 times in PBS, prior to resuspending the cells at a concentration of 1.5E5/mL. For 96-well plate differentiation, cells were resuspended in 200 µL of complete RPMI (cRPMI) + 1.25%v/v mSCF + 10 ng/mL IL-3 (PeproTech Cat# 213-13) + 10 ng/mL Granulocyte Colony Stimulating Factor (GCSF) (PeproTech Cat#300-23) and incubated for 48 hours at 37°C in 5% CO_2_. After 48 hours, cells were pelleted at 300 x g for 5 min and resuspended in 200 µL of cRPMI + 10 ng/mL GCSF for another 24 hour incubation. For 6 well plate differentiation, 1.5E5/mL of cells were resuspended in 6 mL of cRPMI + 1.25% v/v mSCF + 10 ng/mL IL-3 + 10 ng/mL GCSF and incubated for 48 hours. After 48 hours, cells were spun at 300 x g for 5 minutes and resuspended in 6 mL of cRPMI + 10 ng/mL GCSF.

### Generation of KO-ER-HoxB8 cell lines

To generate knockout (KO)-ER-HoxB8 cell lines (KO-HoxB8), CG12 was seeded into a 96 well plate at 50,000 cells/well. CG12 was then transduced with 30µL of an array of custom sgRNA packaged in lentivirus (From Broad, see Table S1). Transduction was performed by spinfection of lentivirus at 1000 x g for 60 min in the presence of 8 µg/mL polybrene (Sigma Aldrich #107689). After spinfection, polybrene was diluted to a final concentration of 2 µg/mL. After 24 hours, half the medium was exchanged with GMP medium to further dilute the polybrene. To select for transduced Cas9-HoxB8 cells, cells were grown in GMP containing 2 µg/mL puromycin 36-48 hours post-transduction and puromycin was maintained in the medium for at least 72 hours. Cells were then expanded and some were single cell cloned by limited dilutions. KOs were confirmed using Western Blots, Flow Cytometry, or production of ROS.

### Infection of HoxB8-PMNs

After 72 hours of growth in the absence of estrogen, PMNs were washed with PBS. For all washes, cells were pelleted at 300 x g for 5 min at RT. Cells were then incubated at Room Temperature (RT) for 30 min in HBSS^-/-^ (Gibco Cat #14170-112). After 30 min, cells were pelleted at 300 x g for 5 min at RT and resuspended in HBSS^+/+^ (Gibco Cat #14025-092) and rested for 15 min at RT, then shifted to 37⁰C for 15 min. For flow cytometry analysis, cells were seeded onto a 96 well round bottom plate and infected with *Yptb* at an approximate MOI of 30:1 for 1 hour at 37⁰C. After infection, cells were washed in PBS and processed for flow cytometry. For measurement of extracellular ROS, cells were seeded in a 96 well 4HBX plate (Co-star #3922) and infected with *Yptb* at an MOI of 20 for 30 min.

### Flow Cytometry Analysis

After infection, cells were washed in PBS at 4°C, prior to incubation with Fixable Viability Dye APC-eFluor780 (eBiosciences Cat# 65-0865-18) for 30 min on ice. For all washes, cells were pelleted at 300 x g for 5 min at 4°C. Cells were then washed with cold flow cytometry buffer (1% FBS in PBS) and blocked with mouse Fc block (BD Pharm Cat# 553142) for 10 min on ice according to manufacturer’s protocol. Cells were then stained with antibodies for 30 min on ice: anti-CD11b-Pacific Blue (Biolegend Cat# 101224), anti-Gr-1 APC (Biolegend Cat# 108412), anti-CD63 PE (BD Pharm Cat# 564222), and CD62L Brilliant Violet 605 (Biolegend Cat#104438). After staining, cells were washed once and fixed in 4% paraformaldehyde for 10 min at RT. After fixation, cells were washed in PBS and stored in flow cytometry buffer until analysis.

When screening cells in a 96-well format for flow cytometry (Fig. 3-4; Fig. S2), cells were analyzed using the BD Celesta (Fig. S2) or Cytek Aurora (Fig. 3-4) and analyzed with FlowJo Software (Version 10.10.0). For CD63 analysis, cells were gated as shown in Fig. S2. To compare the impact of specific Yops in our KO cell lines, we adapted an analysis performed in other published studies on gene interaction mapping and tailored it to our system [59, 60]. For all cell lines tested, the percentage of cells with CD63 localized to the plasma membrane after infection with a Yop-only strain was divided by the plasma membrane localized CD63 after infection with the Δ5-*Yptb* strain, notated as CD63 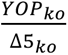. For each KO cell line, this ratio was then divided by the value obtained from the parental CG12 cell line after infection with the same strains on the same day (Fig. 4A). Relative Yop Potency is plotted and calculated as 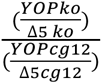

### Total levels of CD63 protein expression

Single cell clones of each cell line were differentiated into Hoxb8-PMNs as described above. Cells were then permeabilized with 0.1% Triton X-100 prior to staining with PE-conjugated anti-CD63 and analyzed on Cytek Aurora Spectral Analyzer on the same day. Cells were gated as shown on Fig. S2 and CD63 gMFI was obtained for each KO cell line and divided by the gMFI of permeabilized CG12.

### Extracellular ROS using isoluminol

ROS isoluminol assays were performed as described previously [4, 40, 61] with the following modifications. Briefly, 4HBX white 96-well (Co-star #3922) plates were incubated and pre-coated with 10% FBS at 37⁰C for at least 1 hour. Plates were washed 3 times with PBS prior to plating cells. Neutrophils were processed for infection as described above and after incubating HoxB8-PMNs in HBSS+/+ for 15 at 37°C, cells were loaded with 25 µM isoluminol (Sigma A8264) and 50 µg/mL HRP (Sigma P6782) and plated onto the pre-coated 4HBX 96 well plates. To synchronize infections, bacteria were spun onto the cells at 209 x g for 1 minute at RT. Chemiluminescence was measured using a BioTek Synergy H1 microplate reader set to 37⁰C with 5% CO_2_ with readings taken every 2 min. To account for variations in cell density and viability between different cell lines, Promega Cell Titer Glo 2.0 (Cat# G9241) was used. When plating cells to measure ROS, cells were also plated in a separate well and mixed with a 1:1 ratio of Cell Titer Glo 2.0, where luminescence values of KO cell lines were divided by values from CG12. This ratio was then multiplied by the Total Chemiluminescence (AUC) from each condition.

### Western Blot

Cells were seeded at 1.5E6 cells per well on a 10% FBS coated 24-well flat bottom plate (non-TC coated) and infected with an MOI of 20 for 5 minutes at 37⁰C 5% CO_2_. After infection, cells were lysed with pre-heated 1x Novex lysis buffer (4X stock: 40% sucrose, 6.82% Tris-Base, 6.66% Tris-HCL, 8% SDS, 0.06% EDTA, 0.075% Bromophenol Blue, 2.5 mM NaVO4,100 mM DTT) and homogenized with pre-chilled Qiagen Qiashredders (Cat# 79656) at max speed in a microcentrifuge for 30 seconds at 4⁰C. Lysates were boiled for 10 min, and 2E5 cells per lane were resolved on a 4-12% Bis-Tris SDS-PAGE (Invitrogen #NP0322BOX).

Samples were immunoblotted with anti-phospho-p44/42 MAPK (ERK1/2) (Thr202/Tyr204) (CST Cat# 9101s; 1:500 dilution), anti-ERK1/2 (CST Cat#9102s; 1:500), or RhoGDI (CST Cat#2564; 1:1000 dilution). Donkey anti-rabbit IRDye 800CW (Cat# 926-32213) secondary antibody was used at 1:20,000. Blots were imaged on Odyssey CLx Li-Cor system and IS Image Studio was used for analysis and quantification.

### Measuring release of granules using ELISA

To assess the amount of granule proteins released, infections were performed as for flow cytometry analyses but supernatants were collected after 1 hour infection and stored at -80⁰C with Halt™ Protease and Phosphatase Inhibitors (Thermofisher Cat. No. 78442) until analysis. After thawing on ice, supernatants were diluted 1:30-1:50 in 1% BSA/PBS. Myeloperoxidase (MPO) was measured using MPO DuoSet ELISA (R&D Systems; Cat #DY3667) according to the manufacturer’s protocol and quantified using a standard curve.

### Mouse Infections

Female BALB/c 8 to 11 week old mice (Charles River Laboratory) or *Skap2^-/-^* [4, 62] were orally infected with a 1:1 mixture of WT-*Yptb*^kan^ and *Yptb* Δ*yopE* via oral gavage with a total of 2E9 CFU in 100ul PBS or intravenously infected with a total of 1000 CFU in 100ul PBS. Three days post-infection, mice were euthanized with CO_2_ and tissues were harvested, homogenized, and outputs were plated on L plate with 0.5 µg/mL irgasan. Outputs were then patched on L^kan^ to determine competitive index by calculating a ratio, normalized to the input:

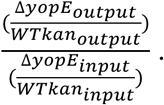

### Bacterial Strains and Growth Conditions

*Yptb* strains used are listed in Table 1. Strains constructed in this study were generated using allelic exchange as previously described [63, 64]. Briefly, the Yop-only strains were constructed using previously published pCVD442 deletion constructs for *yopM* and *yopO* [64, 65]. Bacterial strains were cultured from a single colony plated on L irgasan plates that was grown overnight in 2xYT broth for 16-18 hours at 26⁰C with aeration. For mouse infections, overnight cultures were used for infection and kanamycin^r^ strains were grown in media supplemented with 50 µg/mL kanamycin. For infections of neutrophils, overnight cultures were diluted 1:40 in 2xYT supplemented with 20 mM MgCl_2_ and 20 mM sodium-oxalate. Cultures were incubated at 26⁰C for 1.5 hours with aeration and shifted to 37⁰C for another 1.5 hours with aeration. Bacteria were infected at the MOI indicated in specific experiments and in figure legends.

**Table 1:** *Yersinia* strains used in this study.

| Original Strain # | Description | Reference |
| --- | --- | --- |
| JM301 | YPIII pIB1 | [66] |
| LL167 | YPIII pIB1<br>$\Delta yopEMOJH$ ($\Delta 5$) | [64] |
| ACF123 | YPIII pIB1<br>$\Delta yopEMOJ$ (“H-only”) | This study |
| LL162 | YPIII pIB1<br>$\Delta yopMOJH$ (“E-only”) | This study |
| MM48 | YPIII pIB1<br>$\Delta yopEMJH$ (“O-only”) | This study |
| LL34 | YPIII pIB1 $\Delta yopO$ | [64] |
| ACF125 | YPIII pIB1 $\Delta yopE$ | [64] |
| ACF107 | YPIII pIB1 $\Delta yopH$ | [64] |
| ACF108 | YPIII pIB1 $\Delta yopEH$ | [64] |
| LL45 | YPIII pIB1 $\Delta yopEO$ | [64] |
| LL57 | YPIII pIB1 $\Delta yopHO$ | [64] |
| LL56 | YPIII pIB1<br>$\Delta yopEHO$ | [64] |
| JM495 | YPIII kanR | [67] |
| WS125c | YPIII $\Delta yopE$ | [20] |

### Statistical Analyses

Statistical analysis was performed in GraphPad Prism Version 10 and statistical tests used are indicated in the figure legends.

## Acknowledgements

We thank Dr. Rachel Ende and current members of the Mecsas Lab for critical reading of the manuscript. This work was supported by R01AI169786 from the NIH awarded to JM. PAL was supported by training grant T32AI007422 from the NIH, CMM and EL were supported by training grant T32 GM139772 from the NIH.

