## Supplemental figures for "Neutrophil degranulation and extracellular ROS production are inactivated by *Yersinia pseudotuberculosis* YopE through a SKAP2 independent pathway"

**Supplemental Figure Legends**

**Supplemental Figure 1: YopK is not necessary for blocking CD63 plasma membrane localization** CG12-PMNs were infected with the indicated strains of *Yptb* at an MOI of 30 for 1 hour at 37°C. Δ*yopK* is a strain of *Yptb* lacking YopK. Δ*5* is a strain of *Yptb* lacking the 5 Yop effectors: YopE, YopH, YopO, YopJ, and YopM; Δ6 is a strain of *Yptb* lacking 6 Yop effectors: YopE, YopH, YopO, YopJ, YopM, and YopK. After infection, cells were stained, fixed, and analyzed on Aurora Cytek Spectral Cytometer. Gating Strategy is shown in SF2. Percentage of cells with CD63 localized to the plasma membrane are plotted. Statistical significance was calculated using One-Way ANOVA followed by Sidak's multiple comparison test.

**Supplementary Figure 2: Gating Strategy for assessing CD63 plasma membrane localization** Debris was gated out using FSC-A v. SSC-A, followed by gating for single cells using FSC-A v. FSC-H. Single Cells were analyzed with a Viability Dye. Viability Dye negative single cells were subsequently analyzed for CD11b and Gr-1 expression. CD11b+Gr-1+ population was further analyzed for percentage of CD63+ in uninfected cells and cells infected with *Yptb*-Δ*5* plotted as a histogram. For some analysis, CD63 geometric Mean Fluorescence Intensity (gMFI) was used and obtained from Live Singlets expressing CD11b+ and Gr-1+.

**Supplemental Figure 3: Preliminary Screen with pooled knockout cell lines** (A) Histogram of surface localized CD63 on CG12-PMNs (black, dashed), two CD63-KO-PMN pools generated with two different gRNAs (blue), and unstained CG12-PMN samples (grey, dashed). (B-C) Each well of a 96-well plate containing CG12 were transduced with a gRNA targeting the indicated gene on the x-axis. After 9 days of 2 µg/mL puromycin selection, 2.5-3 x $10^4$ cells per well were seeded into two 96-well plates and differentiated for 3 days. Cells were (B) infected with *Yptb*-Δ*5* at an MOI of 30 or (C) left uninfected for 1 hour, and processed for CD63 plasma membrane expression. Cells were analyzed on BD FACSCelesta. (B) CD63 gMFI of cell lines infected with *Yptb*-Δ5; values 1 standard deviation above (green circles) or below (blue circles) are indicated. Shaded regions represent mean ± 1 SD, with mean being marked with a solid line. n=1 (C) Normalized CD63 gMFI relative to parental CG12 (set at 1) for uninfected parental pools of cell lines that were single cell cloned. NTC, non-targeting control using a gRNA targeting luciferase.

**Supplemental Figure 4: Validation of KO in single cell clones** Lysates from single cell clones and CG12-PMN were resolved on a Bis-Tris Gel and probed with antibodies for the indicated protein. Confirmed single cell clones used in the screen were denoted with an asterisk (*). (A) Lysates from PLCγ2-KO-Hoxb8 GMP single cell clones and CG12-PMN were probed with an α-PLCγ2 antibody Cell Signaling Cat#3872S; (B) Lysates from SYK-KO PMN single cell clones and CG12 were probed with an α-SYK antibody (Cat# 2712s); (C) Lysates from SLP76-KO PMN single cell clones and CG12 were probed with an α-SLP6 antibody (Cat# 05-1426); (D) Lysates from Pyk2-KO PMN single cell clones and CG12 were probed with an α-Pyk2 CST 3292s; (E) Lysates from LYN-KO PMN single cell clones and CG12 were resolved on a Bis-Tris Gel and probed with an α-Lyn Ab Cat# MA1-19334; (F) Lysates from VAV1-KO GMP single cell clones and CG12 were probed with an α-Vav1 monoclonal Ab (Upstate Cat 07-192; lot 23747); (G) Lysates from MPO-KO-Hoxb8 GMP single cell clones and CG12-PMN were probed with an α-MPO antibody AbCam Cat#9535; (H) Lysates from BTK-KO GMP and CG12-PMN were probed with an α-BTK antibody (CST #8547S); (I) Lysates from SKAP2-KO-PMN and CG12-PMN were resolved on a Bis-Tris Gel and probed with an α-SKAP2 antibody (Proteinech Cat# 12926-1-AP); (A-I) RhoGDI was used as loading control (Cell Signaling Cat#2564S). (J) RhoG-GMP single cell clones were probed with an α-RhoG antibody (SC-80015). Beta-actin was used as loading control (SC-4778); (K) PXN-KO-PMN single cell clones were probed with an α-PXN antibody (BD Pharm 610051).

**Supplemental Figure 5: Validation of eGFP-KO and CYBB-KO clones** (A) CG12, WT-ER-HoxB8, and GFP-KO-GMP clones from sgRNA Well H9 subclones #1-5 were analyzed on LSRII flow cytometer for GFP expression. Subclone eGFP-KO H9-3 was used for screen*. (B) CG12, *Cybb-/-* PMN, and CYBB-KO clones from sgRNA Well B2 subclones #1-10 were differentiated into PMNs as previously described and assessed for extracellular ROS production after PMA stimulation. Subclone CYBB-KO B2-8 was used for screen*

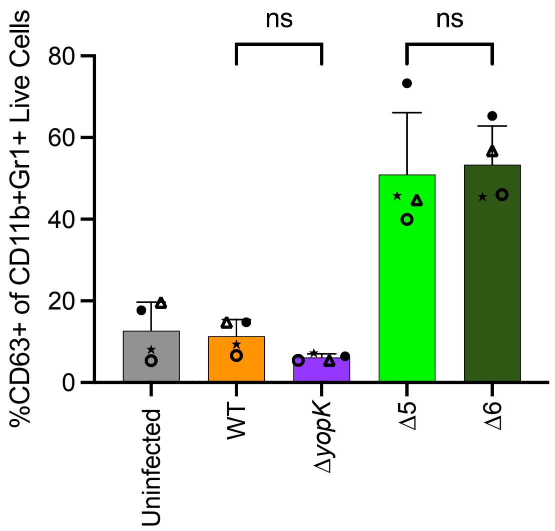


**Supplemental Figure 1**

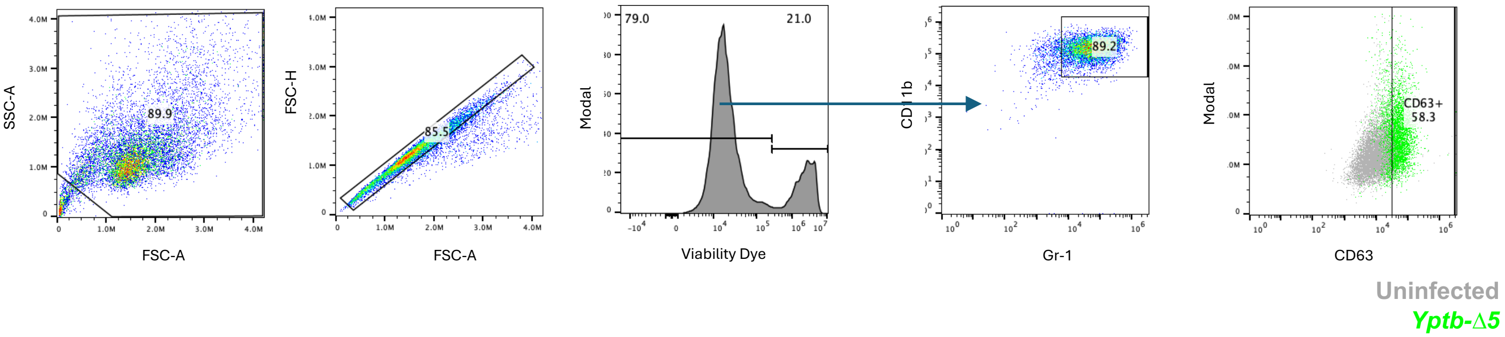


**Supplementary Figure 2**

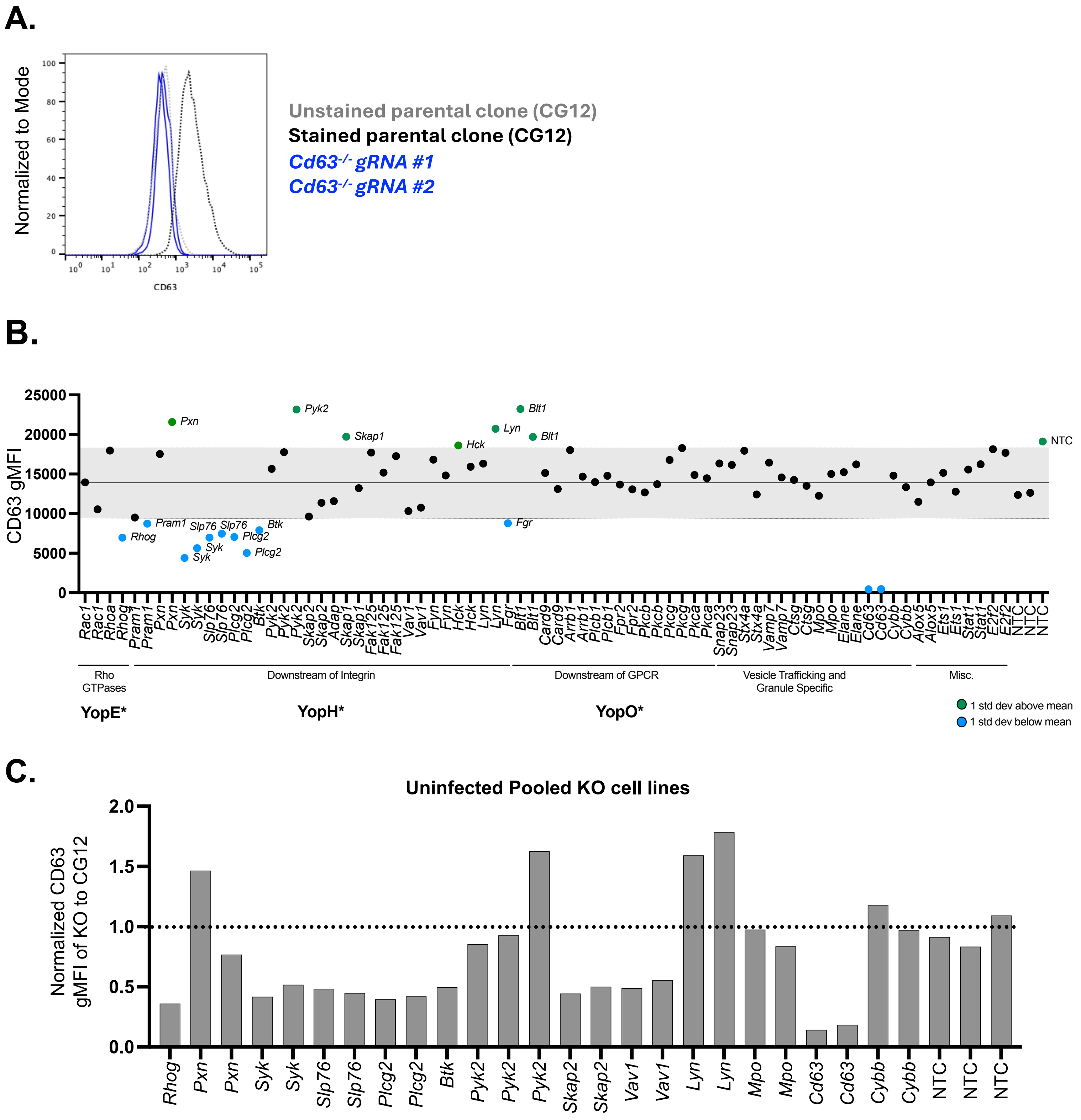


Supplemental Figure 3

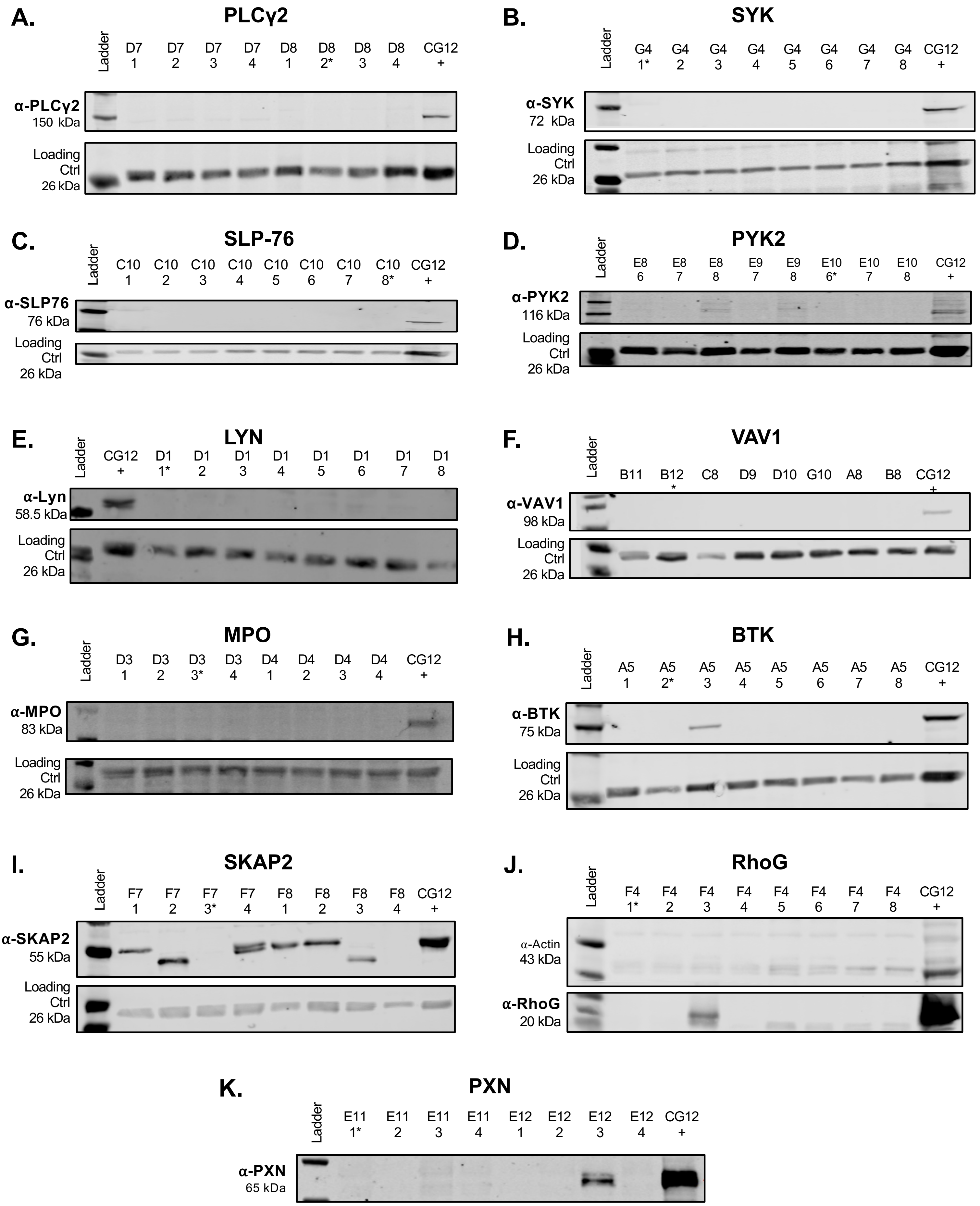


**Supplemental Figure 4**

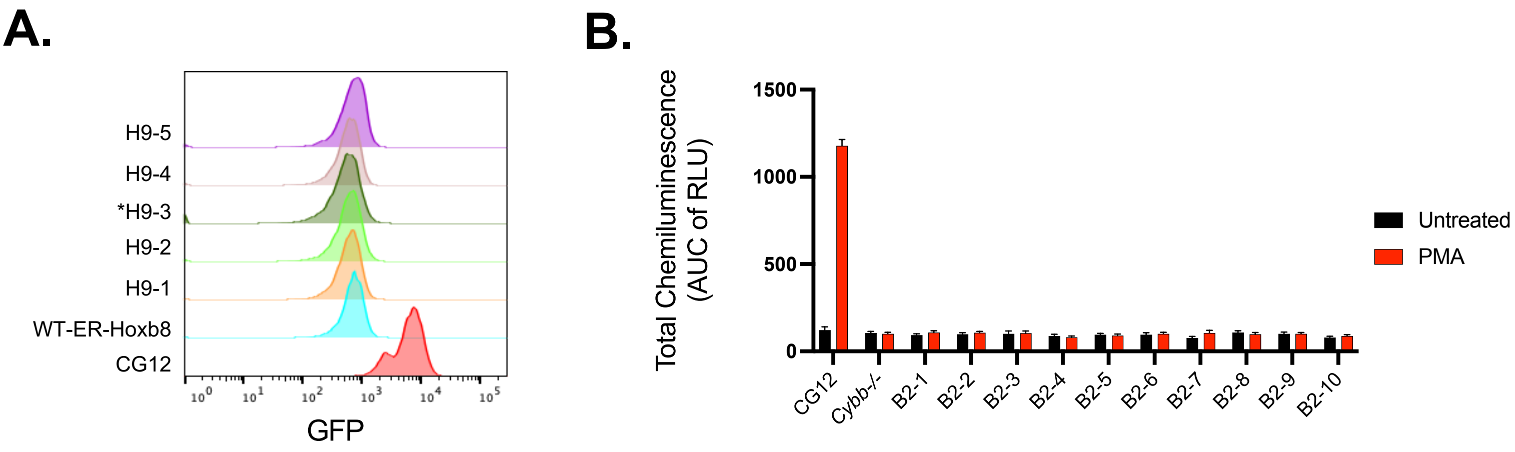


**Supplemental Figure 5**
